# SEVI Fibrils are Induced by Bacterial Surface Molecules and Exert Antimicrobial Activity Against ESKAPE Pathogens

**DOI:** 10.1101/2025.01.31.635909

**Authors:** Verena Vogel, Lia-Raluca Olari, Richard Bauer, Sandra Burczyk, Fiona Witz, Emely Walker, Nourice Jaber, Stefanie Mauerer, Ulrich Rupp, Ulrich Stifel, Julia Zinngrebe, Clarissa Read, Pamela Fischer-Posovszky, Jan Münch, Barbara Spellerberg

**Author notes:** Corresponding authors: Jan Münch and Barbara Spellerberg.

## Abstract

The semen-derived enhancer of viral infection (SEVI) is an amyloid fibril formed by the self-assembly of the PAP248-286 peptide, a cleavage product of prostatic acid phosphatase, which is naturally present in human seminal plasma. Beyond its previously described role in HIV transmission, findings highlight a physiological function for SEVI in innate immunity. As many amyloids display antimicrobial effects, we tested SEVI fibrils for antibacterial activity against microbial ESKAPE pathogens and urogenital bacteria, including *Pseudomonas aeruginosa, Klebsiella quasipneumoniae, Escherichia coli, Acinetobacter baumannii, Staphylococcus aureus, Streptococcus agalactiae* and *Listeria monocytogenes*. SEVI exhibited direct dose-dependent antibacterial effects in radial diffusion and survival assays, with activity observed at physiologically relevant concentrations. Bacterial surface molecules such as lipopolysaccharides and lipoteichoic acid induced the formation of SEVI fibrils, as confirmed by kinetic assays. Preincubation with epigallocatechin gallate, a fibril disruptor, abolished SEVI’s antibacterial activity, pointing to the importance of its fibrillar structure. Mechanistic studies and electron microscopy revealed limited bacterial membrane disruption and the intracellular accumulation of polyphosphate granules in *P. aeruginosa*, indicating a stress response. In conclusion, SEVI exerts a potent antibacterial activity against pathogens found in the urogenital tract, indicating a potential physiological role in vaginal mucosal immunity.

## Introduction

Historically, amyloids have been studied primarily in the context of degenerative diseases, such as Alzheimer’s and prion diseases. However, more recently their antimicrobial properties came into focus^1^. Several amyloids and amyloidogenic peptides, including Aβ from Alzheimer’s disease^2^, prion proteins^3^, serum amyloid A^4^, β2-microglobulin^5^, alpha-synuclein^6^ and hemoglobin-derived peptides^7^, display antibacterial activity. These amyloids destroy bacteria by disrupting bacterial membranes or promoting aggregation, which facilitates their phagocytotic clearance. Interestingly, some well-characterized antimicrobial peptides (AMPs), including LL-37^8^, lactoferrin^9^, and lysozyme^10^, also form amyloid structures, hinting at an evolutionary link between amyloid formation and antimicrobial function. ^11^

Semen-derived enhancer of viral infection (SEVI) was first identified as a fibrillar amyloidogenic peptide generated through the proteolytic cleavage of amino acids 248– 286 of prostatic acid phosphatase (PAP). SEVI fibrils significantly enhance HIV-1 infection by facilitating the binding and fusion of viral particles with the negatively charged membranes of HIV-1 virions and host cells ^12^. This effect is attributed to the highly positive charge of SEVI fibrils at neutral pH values. While these fibrils promote HIV-1 infection, their primary biological function is likely to facilitate removal of damaged sperm and bacterial clearance. Semen amyloids have been shown to immobilize sperm, enabling the clearance of damaged and apoptotic cells via macrophages in the female reproductive tract^13^. This process may contribute to reproductive health by aiding in sperm selection and antigen removal. Additionally, SEVI both interacts with both Gram-positive and Gram-negative bacteria in a charge-dependent manner, promoting bacterial aggregation and phagocytosis by macrophages ^14^. In summary, these findings indicate that semen amyloids may play an important role in reproduction and innate immune mechanisms.

PAP248–286 exhibits AMP-like characteristics^15^, such as a positive charge and amphipathic structure, enabling strong interactions with negatively charged bacterial surfaces. Therefore, we investigated the freshly dissolved non-fibrillar PAP248–286 peptide and its fibrillar form (SEVI) for their antibacterial activity against a range of bacterial pathogens, including the ESKAPE species *Enterococcus faecium, S. aureus, K. quasipneumoniae, A. baumannii*, and *P. aeruginosa*. These pathogens are often associated with multidrug resistance, and are major contributors to life-threatening nosocomial infections, particularly among critically ill and immunocompromised patients. Together, they account for a significant portion of the five million annual deaths attributed to antimicrobial resistance^16^. All ESKAPE pathogens are categorized as critical or high-priority threats on the 2024 WHO bacterial priority pathogens list, emphasizing the urgent need for innovative antimicrobial strategies ^17^. Identifying agents capable of targeting these pathogens is essential for addressing severe bacterial infections and advancing our understanding of innate immune mechanisms. Beyond ESKAPE pathogens, we also examined SEVI’s effects against *S. agalactiae* and *L. monocytogenes*, both of which are key players in urogenital and pregnancy-associated infections.

Given SEVI’s ability to facilitate bacterial uptake by phagocytes^14^, we hypothesize that SEVI may function as a natural antimicrobial agent, contributing to immune defense in mucosal environments such as the urogenital tract. Thus, we here explored the antimicrobial properties of non-fibrillar and fibrillar PAP248–286 (SEVI) and characterized the associated antibacterial mechanisms, which led to the detection of intracellular phosphate bodies and observation of accelerated SEVI fibril formation upon contact with bacterial surface molecules. With the growing prevalence of multidrug-resistant pathogens, this study aims to provide new insights into the dual role of amyloidogenic peptides in pathogen defense and their potential as therapeutic agents against resistant bacterial infections.

## Material and Methods

### Culture conditions

All bacterial strains used in this study are listed in Table 1. For liquid culture, *E. coli, P. aeruginosa*, and *A. baumannii* were grown in lysogeny broth (LB-Miller) at 37°C with shaking (160 rpm) under aerobic conditions. *L. monocytogenes* was cultivated in brain-heart-infusion medium (Oxoid). All other bacteria were cultured in THY medium (Todd-Hewitt Broth [Oxoid] supplemented with 0.5% yeast extract [BD, Miami, FL]) at 37°C in a 5% CO_2_ atmosphere. For plate cultivation, tryptone soy agar supplemented with 5% sheep blood (Oxoid) was used, with incubation overnight at 37°C in 5% CO_2_. *L. monocytogenes* ^18^ and *S. agalactiae* transformed with pNZ-pHin2LM ^19^ were grown in medium containing chloramphenicol at 10 or 6 µg/ml (Sigma-Aldrich), respectively. Additionally, *L. monocytogenes* pNZ-pHin2LM ^19^ was grown aerobically in brain-heart-infusion medium (Oxoid) at 37°C with shaking (160 rpm), while *S. agalactiae* was grown statically in THY medium.

**Table 1:**
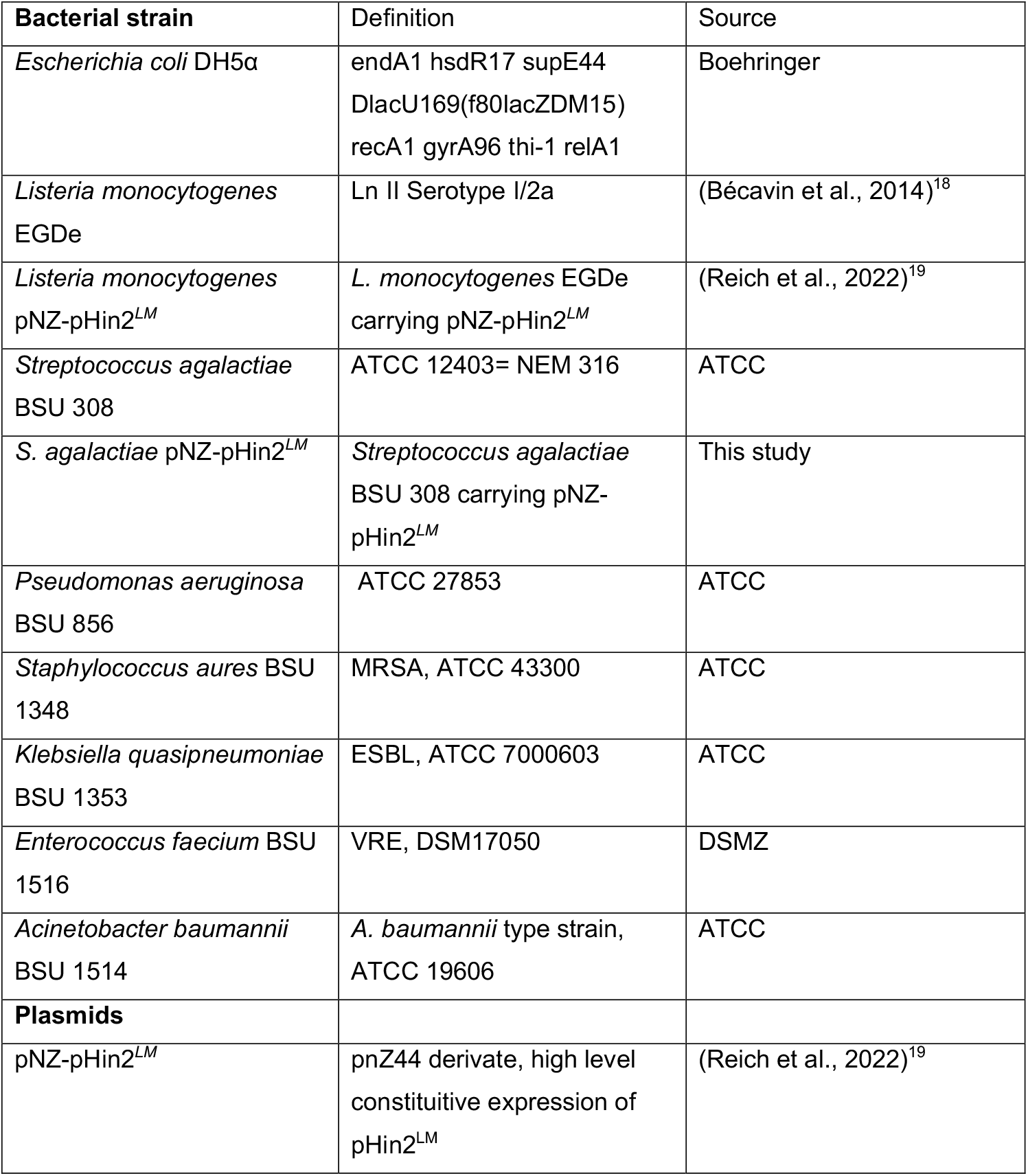
Bacterial strains and Plasmids used in this study.

| Bacterial strain | Definition | Source |
| --- | --- | --- |
| *Escherichia coli* DH5α | endA1 hsdR17 supE44<br>DlacU169(f80lacZDM15)<br>recA1 gyrA96 thi-1 relA1 | Boehringer |
| *Listeria monocytogenes*<br>EGDe | Ln II Serotype I/2a | (Bécavin et al., 2014)<sup>18</sup> |
| *Listeria monocytogenes*<br>pNZ-pHin2<sup>LM</sup> | *L. monocytogenes* EGDe<br>carrying pNZ-pHin2<sup>LM</sup> | (Reich et al., 2022)<sup>19</sup> |
| *Streptococcus agalactiae*<br>BSU 308 | ATCC 12403= NEM 316 | ATCC |
| *S. agalactiae* pNZ-pHin2<sup>LM</sup> | *Streptococcus agalactiae*<br>BSU 308 carrying pNZ-pHin2<sup>LM</sup> | This study |
| *Pseudomonas aeruginosa*<br>BSU 856 | ATCC 27853 | ATCC |
| *Staphylococcus aureus* BSU<br>1348 | MRSA, ATCC 43300 | ATCC |
| *Klebsiella quasipneumoniae*<br>BSU 1353 | ESBL, ATCC 7000603 | ATCC |
| *Enterococcus faecium* BSU<br>1516 | VRE, DSM17050 | DSMZ |
| *Acinetobacter baumannii*<br>BSU 1514 | *A. baumannii* type strain,<br>ATCC 19606 | ATCC |
| **Plasmids** |  |  |
| pNZ-pHin2<sup>LM</sup> | pnZ44 derivate, high level<br>constitutive expression of<br>pHin2<sup>LM</sup> | (Reich et al., 2022)<sup>19</sup> |

### Radial Diffusion Assay

The antimicrobial activity of SEVI was evaluated using a radial diffusion assay (RDA)^20^. Overnight cultures of target bacteria were centrifuged at 3000 x g for 10 minutes, washed with 10 mM phosphate buffer, and centrifuged again. The pellet was resuspended in 10 mM phosphate buffer, and the optical density at 600 nm (O.D. 600) was measured. Approximately 2 x 10^7^ bacterial cells were then transferred into 15 ml of 1% agarose. After pouring the agar plates and allowing them to dry for 30 minutes at 4°C, wells were created using sterile wide-bore pipette tips (Axygene, a Corning brand). Each well was filled with PAP248-286 (Celtek Peptides) or its fibrillar form, SEVI (Semen-derived Enhancer of Viral Infection) that were formed by shaking the dissolved peptide for 7 days at 1,500 × rpm, 37°C. Plates were incubated at 37°C for 3 hours, followed by an overlay with 10 ml of trypticase soy agar (Oxoid). After overnight incubation at 37°C in 5% CO_2_, inhibition zones, visible as clear halos around the wells, were measured. To assess the effect of pH on the antimicrobial efficacy of both the peptide and fibril forms, the agarose and overlay agar were adjusted to various pH levels (5, 5.5, 6, 6.5, and 7) using NaOH and HCl prior to performing the RDA. Additionally, SEVI was preincubated with (-)-epigallocatechin-3-gallate (EGCG) (Sigma) for 3, 22, or 45 hours to determine the role of its amyloid structure in antibacterial activity, as EGCG is known to disaggregate fibrils ^21^. This pretreatment allowed for analysis of whether antibacterial effects were mediated by the fibrillar or monomeric form of SEVI. To determine whether the fibrillar or non-fibrillar form of PAP 248-286 exhibits antibacterial activity, non-fibrillar PAP248-286was agitated for 17–18 hours. Following this, the sample was centrifuged at 13,300 × g for 10 minutes. The resulting supernatant, containing non-fibrillar PAP248-286, and the pellet, containing the fibrils, were then separately tested for their inhibitory activity in RDA.

### Bacterial Survival Assay

The antimicrobial activity of monomeric or fibrillar PAP248-286 in a liquid environment was assessed using a survival assay. A 1 ml aliquot of an overnight bacterial culture, diluted to an O.D. 600 nm of 0.1, was centrifuged at 17,000 x g for one minute. The resulting bacterial pellet of *P. aeruginosa* was resuspended in 20 mM phosphate buffer adjusted to a pH of 4.5. The pellet of *L. monocytogenes* was reconstituted in vaginal fluid simulant (VFS) (containing 24.7 mM glucose, 77 mM NaCl, 3.78 mM sodium acetate, 18 mM sodium lactate, 6.48 mM L-lactate, and 0.79 ml glacial acetic acid in 1 liter of water; pH 4.2). Subsequently, bacterial suspensions were mixed with 50 µg/ml of PAP248-286 or SEVI, water was used as the untreated control. Incubation at 37°C for 2 hours was performed, samples were taken at 0, 1, and 2 hours, serially diluted, and plated on agar. After overnight incubation at 37°C in 5% CO_2_, colonies were counted, and colony-forming units (CFU) per ml were determined.

### SYTOX Assay

To investigate whether SEVI has an effect on the bacteria membrane a SYTOX Green Membrane Permeabilization Assay was performed with *L. monocytogenes* and *P. aeruginosa*, as previously described ^22 20^. In short, bacteria were freshly inoculated and grown till an O.D. 600 nm of 0.1 was reached. After centrifugation the pellet of 1 ml was solved in 20 mM sodium phosphate buffer with 10% trypticase soy broth (Oxoid) and 0.2 μM SYTOX (Invitrogen), adjusted to a pH of 7 or 4.5. 90 µl of bacteria mixed with SYTOX were filled in a 96-well plate and treated with 10 µl of different PAP248-286 concentrations. End concentrations 100 and 3.12 µg/ml. Fluorescence emission at 530 nm was measured after excitation at 488 nm with a Tecan microplate reader M infinite 200 reader (Tecan group). Bacterial cells treated with 70% ethanol were used as a positive control and water treatment as a negative control. Measurements were performed in triplicates and three independent experiments were conducted.

### Generation of *S. agalactiae* carrying a pHluorin plasmid

Electroporation of *S. agalactiae* was performed as previously described ^23^. In brief, 100 ml of a dense *S. agalactiae* culture were washed multiple times with 10% glycerol and finally resuspended in 20 % glycerol. Bacterial cells were exposed to an electrical pulse in the presence of 1 µg pNZ-pHin2^*LM*^. After a 2-hour incubation period at 37°C, the cells were plated on THY agar supplemented with 6 µg/mL chloramphenicol (Sigma-Aldrich). The resulting colonies were examined for plasmid presence and a fluorescence signal.

### pHluorin assay

To further examine the effect of SEVI on bacterial membrane integrity, a pHluorin assay was performed, as previously described^19^. In brief, an overnight culture of *L. monocytogenes* pNZ-pHin2LM or *S. agalactiae* pNZ-pHin2LM was adjusted to an O.D. 600 nm of 3.0 in Listeria minimal buffer (200 mM MOPS, 4.82 mM KH_2_PO_4_, 11.55 mM Na_2_HPO_4_, 1.7 mM MgSO_4_, 0.6 mg/ml (NH_4_)_2_SO_4_, 55 mM glucose pH 6.2) ^24^. 50 µl aliquots of bacteria were mixed with 50 µl of Listeria minimal buffer supplemented with SEVI in concentrations ranging from 100 to 0.1 µg/ml. After incubation for 30 min at 37°C in the dark, fluorescence intensity was measured with a Tecan microplate reader M infinite 200 (Tecan group). Emission was measured at 520 nm after excitation at 400 and 480 nm. Listeria minimal buffer alone served as negative control and 100 µg/ml nisin (Sigma-Aldrich) as positive control.

### Electron microscopy

To assess pore formation in bacterial membranes, transmission electron microscopy (TEM) was used to analyze the effects of PAP248-286 and SEVI on *S. agalactiae* and *P. aeruginosa*. Overnight bacterial cultures were freshly inoculated at an initial O.D. 600 nm of 0.02 and grown to an O.D. 600 nm of 0.1. A 1 ml aliquot of each culture was centrifuged at 8800 x g for 2 minutes, and the resulting pellet was resuspended in 90 µl of 20 mM phosphate buffer (pH 4.5) containing 10% tryptic soy broth (TSB). Subsequently, 10 µl of PAP248-286 or SEVI was added to achieve a final concentration of 50 µg/ml, followed by incubation at 37°C for 60 minutes. As a negative control, bacterial cells were incubated with water.

After incubation, samples were centrifuged again at 8800 x g for 2 minutes, and 60 µl of the supernatant was removed. The remaining 40 µl of bacterial suspension was mixed with 40 µl of double-concentrated fixative solution (5 % glutaraldehyde, 1 % sucrose in phosphate buffer). Samples were then processed via postfixation with osmium tetroxide, dehydrated in a graded series of propanol, embedded in Epon resin, block contrasted with uranyl acetate, ultrathin-sectioned and contrasted with lead citrate as described previously ^7 10^. Imaging was conducted using a Joel 1400 transmission electron microscope at 120 kV acceleration voltage. Additionally, energy-dispersive X-ray spectroscopy (EDX) was performed to analyze the elemental composition of granules and cytoplasm in *P. aeruginosa* samples, using a Hitachi S-5200 field emission scanning electron microscope equipped with a scanning transmission electron microscopy (STEM)-detector (Hitachi High-Tech Corp., Tokyo, Japan) and an EDAX Phoenix X-ray detector system (AMETEK GmbH, Weiterstadt, Germany). Acceleration voltage for EDX was 30 kV and section thickness was 30 nm. A minimum of 20 images per sample was captured for analysis.

### Thioflavin T assay

To determine whether LPS or Lipoteichoic Acid (LTA) could similarly induce fibril formation of PAP248-286, Thioflavin T assays were carried out. A Thioflavin T (ThT) (Sigma-Aldrich) stock solution of 2.5 mM was prepared in PBS and sterile-filtered. PAP248-286 monomers at a concentration of 400 µg/ml were mixed with different concentrations (0-200 µg/ml) of LPS from *P. aeruginosa* or *E. coli* (Sigma-Aldrich), or Lipoteichoic Acid (LTA) from *Streptococcus pyogenes* (Sigma-Aldrich), and ThT diluted in PBS at a final concentration of 25 µM. Of each preparation, 45 µl were transferred to one well of a Corning® 385-well black polystyrene microplate (Sigma-Aldrich), and two glass beads (1-2 mm diameter) were added to each well. The plate was sealed and incubated at 37°C. Fluorescence was measured every 5 minutes, at an excitation wavelength of 450 nm and an emission endpoint of 490 nm using a Synergy H1 hybrid multi-mode reader (Biotek) and the Gen5 software. Before each measurement, samples were shaken for 30 seconds at 307 cpm (5 mm) to ensure sample homogenization. All values represent fluorescence intensity derived from averaged quadruplicates corrected for the background signal.

### Phagocytic uptake of bacteria

THP-1 monocytes were seeded at 50,000 cells/well into a 96 well plate in RPMI medium supplemented with 10% FCS, 2% L-Glutamine (Gibco), and 55 nM b-mercaptoethanol, and 10 ng/ml PMA (Phorbol-12-myristat-13-acetat, Sigma-Aldrich P1585) was added to induce differentiation into macrophages. After 3 days, cells were washed and fresh medium without FCS and PMA was added. Cells were subsequently treated with different concentrations (2.5 – 50 µg/ml) of PAP248-286 or SEVI as well as pHrodo Red *S. aureus* BioParticles (ThermoFisher A10010) for 3 hours. pHrodo Red *S. aureus* BioParticles show fluorescence when taken up into the phagosomes of macrophages. As negative control, pHrodo Red *S. aureus* BioParticles were added together with 5 µg/ml Cytochalasin D (Sigma-Aldrich C2618). As positive control, we used *S. aureus* BioParticles Opsonizing Reagent (5µl/100µl *S. aureus* BioParticles, ThermoFisher S2860) to opsonize pHrodo Red *S. aureus* BioParticles for one hour at 37°C before adding them to the cells. The plate was imaged in an Incucyte S3 Live Cell Analysis System (Sartorius). Four images were taken in each well every 30 minutes for 3 hours. Analysis of images was automated using the Incucyte 2019B Basic Analysis Software module. A threshold for positive fluorescent signal was applied and the number of positive signals was counted for each well. Each experiment was normalized to the respective controls.

## Results

### Antimicrobial activity of PAP248–286 in its non-fibrillar and fibrillar forms across bacterial species at different pH values

SEVI fibrils have previously been implicated to participate in innate immunity. Enhanced phagocytosis was demonstrated to be mediated by SEVI fibrils^14^ for *S. aureus*. To expand on these findings, we tested the effects of the non-fibrillar PAP248– 286 and SEVI fibrils on phagocytosis of *S. aureus* by THP-1 monocytes. Following incubation with physiological concentrations of the peptide/fibrils, a significantly enhanced phagocytosis activity could be observed for SEVI fibrils after 30 and 60 min, while non-fibrillar PAP248–286 had no effect (Fig. S1), confirming previous data^14^.

To explore whether SEVI fibrils, like other amyloidogenic peptides, exhibit direct antimicrobial activity, we investigated non-fibrillar and fibrillar PAP248–286 against various bacterial species using radial diffusion assays (Fig. 1). These included *E. coli*, the ESKAPE pathogens, *S. agalactiae*, and *L. monocytogenes*. Interestingly, for all bacterial species we tested, the fibrillar form showed equivalent or higher activity than the non-fibrillar preparation. The influence of pH (neutral to acidic) was assessed to reflect the physiological conditions of the urogenital mucosal environment. *P. aeruginosa* demonstrated pronounced susceptibility under acidic conditions. At pH 4.5, inhibition was notably enhanced, with strong effects evident even at lower concentrations. However, *K. quasipneumoniae* showed a contrasting response, with both preparations demonstrating greater inhibition at neutral pH (7.0) compared to acidic conditions (pH 4.5). Overall, no clear trend could be observed in regard to acidic pH. Furthermore, no antibacterial effects could be detected against *S. aureus* and *E. faecium* (Table S1).

**Fig. 1.**
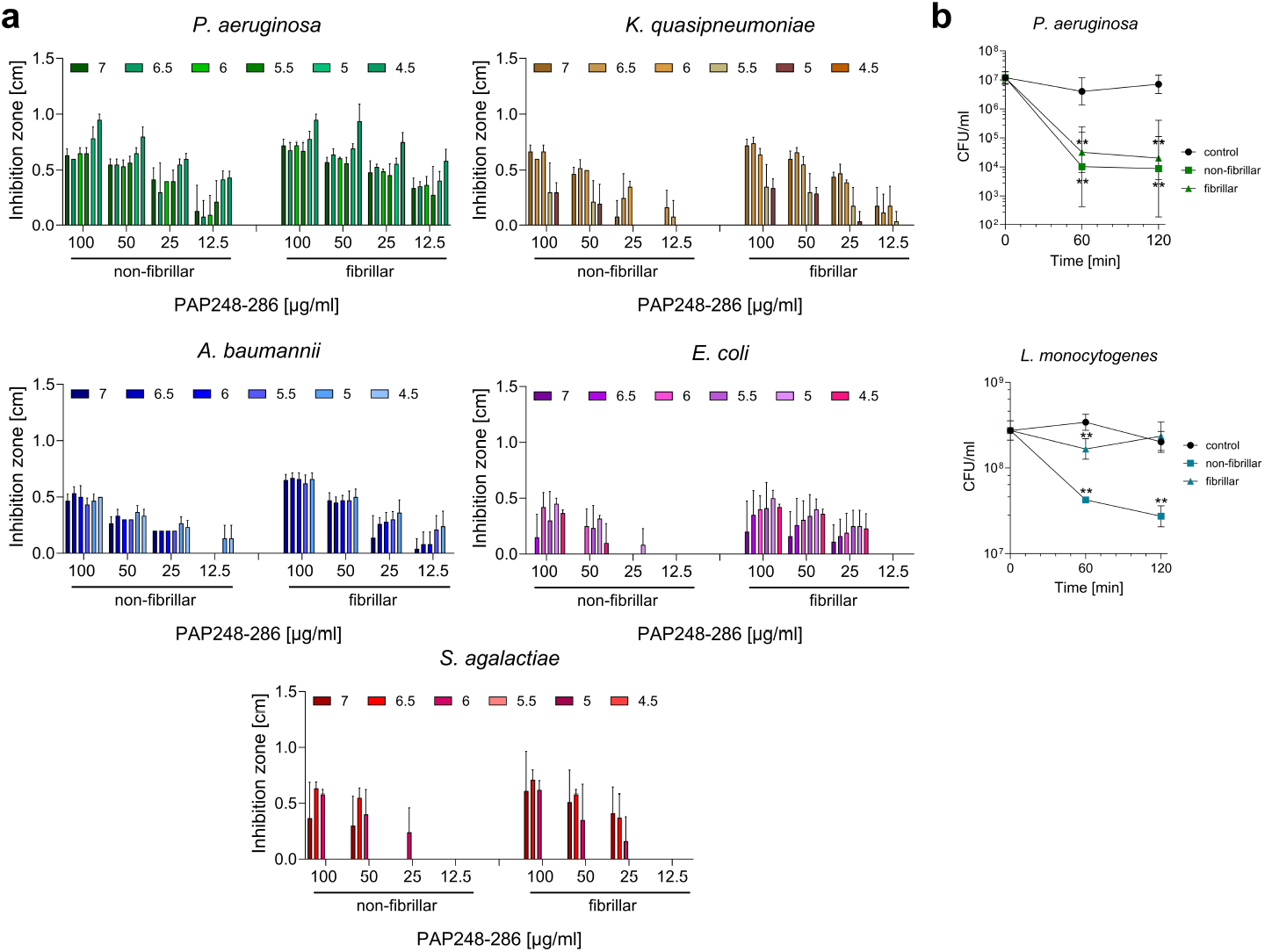
Antibacterial activity of fibrillar and non-fibrillar PAP248-286. a) Fibrillar and non-fibrillar PAP248-286 were tested in a radial diffusion assay against *Pseudomonas aeruginosa, Klebsiella quasipneumoniae, Acinetobacter baumannii, Escherichia coli* and *Streptococcus agalactiae*. The assay was performed at different pH, ranging from 7 to 4.5. b) Survival of *P. aeruginosa* and *Listeria monocytogenes* was monitored over two hours at a pH of 4.5. Bacterial cells were incubated with 50 µg/ml of either fibrillar or non-fibrillar PAP248-286. Colony forming units is abbreviated to CFU. At least five independent experiments were conducted and depicted is the mean ± standard deviation. Figures were created using GraphPad Prism version 10.

Given the antimicrobial activity of non-fibrillar and fibrillar PAP248–286, we further examined its bactericidal effect over 1 and 2 hours using survival assays (Fig. 1b). Incubation with *P. aeruginosa* resulted in a rapid decline in bacterial survival, with the most pronounced reduction occurring within the first 60 minutes, suggesting a fast-acting mechanism that compromises bacterial integrity. The fibrillar as well as the non-fibrillar form show a similarly high antimicrobial activity. In the case of *L. monocytogenes*, non-fibrillar PAP248–286 induced a steady reduction in survival. (Fig. 1b), while SEVI only caused a small reduction in survival after 60 min of incubation.

Taken together, PAP248–286, in both its non-fibrillar and fibrillar (SEVI) forms, exhibits antimicrobial activity with notable species-specific and structural differences: SEVI demonstrates slightly higher, dose-dependent inhibition across different bacterial species and enhanced efficacy at acidic pH for some species, while non-fibrillar PAP248–286 shows time-dependent activity with slower but sustained effects.

### Epigallocatechin-gallate (EGCG) abrogates the antimicrobial activity of SEVI

To investigate whether the fibrillar nature of SEVI is required for its antimicrobial activity, we studied the effects of epigallocatechin-gallate (EGCG), a compound known to disrupt PAP248-286 fibrils ^25^. SEVI was preincubated with EGCG for 3, 22, and 45 hours and subsequently tested for its antimicrobial activity in radial diffusion assays against *P. aeruginosa*. The results demonstrated a time-dependent reduction in the antimicrobial effects of SEVI (Fig. 2). After 3 hours of incubation with EGCG, a partial loss of activity was observed. This loss became more pronounced after 22 hours and reached near-complete abrogation of the antimicrobial activity of SEVI after 45 hours (Fig. 2). These findings suggest that EGCG progressively disassembles SEVI fibrils over time, ultimately eliminating their antibacterial properties. Importantly, EGCG did not affect the activity of LL-37, a known antimicrobial peptide used as a positive control, confirming the specificity of its action on SEVI fibrils. EGCG alone at 1 mM did not show antibacterial effects in RDA testing (Table S2).

**Fig. 2.**
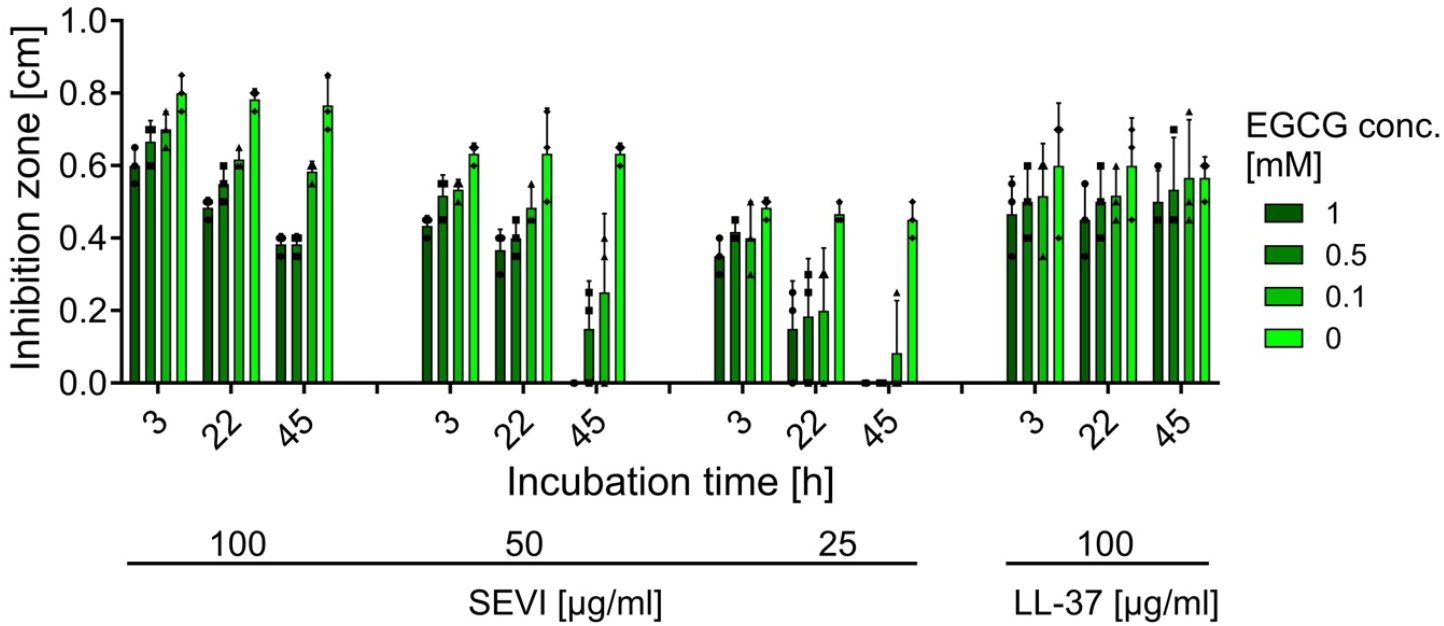
Importance of PAP248-286 fibrils for antimicrobial activity. SEVI concentrations ranging from 100-25 µg/ml were preincubated with various Epigallocatechin gallate (EGCG) concentrations for 3, 22 or 45 h, afterwards the antimicrobial activity against *P. aeruginosa* was assessed in a radial diffusion assay. As a control LL-37 was incubated with EGCG and tested for altered activity. Depicted is the mean + standard deviation of three independent experiments. Figure was created using GrapPad Prism version 10.

To further examine the contributions of SEVI fibrils and non-fibrillar PAP248–286, SEVI fibrils were separated from the supernatant containing monomers and oligomers via centrifugation and tested independently. Radial diffusion assays showed that both the fibrils and the supernatant retained antimicrobial activity against *P. aeruginosa* (Fig. S2). However, preincubation with EGCG primarily disrupted the activity of the fibrillar form, underscoring the importance of SEVI’s structural integrity for full antimicrobial efficacy.

Taken together, the results from different experimental approaches demonstrate that SEVI’s fibrils display antimicrobial activity, which is abrogated by EGCG in a time-dependent manner. Furthermore, the findings suggest that non-fibrillar PAP248–286 retains partial antimicrobial activity, highlighting its potential role as a precursor to fibril formation in bacterial environments.

### LPS and LTA induce PAP248-286 fibrillation

It is well established that negatively charged antimicrobial peptides (AMPs) interact with bacterial cell surface molecules such as lipopolysaccharides (LPS) ^26^, which are found on Gram-negative bacteria, and lipoteichoic acids (LTA), present on Gram-positive bacteria ^27^. These interactions are critical for disrupting bacterial membranes and modulating immune responses. Considering that amyloid formation may be triggered by different molecules including LPS ^28^, we investigated whether LPS derived from *P. aeruginosa* or *E. coli* or LTA derived from *S. pyogenes* affects the formation of SEVI fibrils.

For these experiments, freshly dissolved PAP248-286 was incubated with varying concentrations of LPS, and the formation of SEVI fibrils was monitored over time using a fluorescence-based assay, which provides insights into the aggregation kinetics of the peptide. A concentration of PAP248-286 was chosen that typically does not form fibrils under the tested conditions in the absence of other factors (Fig. 3). However, in the presence of low doses of LPS (10–50 µg/mL) from *P. aeruginosa* (Fig. 3a) or *E. coli (*Fig. 3b), fibril formation was detected. At higher concentrations of LPS (200 µg/mL), this effect was amplified, leading to significantly enhanced fibrillation over time (Fig. 3a and 3b). The fluorescence intensity over time revealed a substantial increase in fibril formation with both *P. aeruginosa* and *E. coli* LPS compared to controls, indicating that LPS accelerates the aggregation of PAP248-286 into amyloid fibrils in in a dose-dependent manner.

**Fig. 3.**
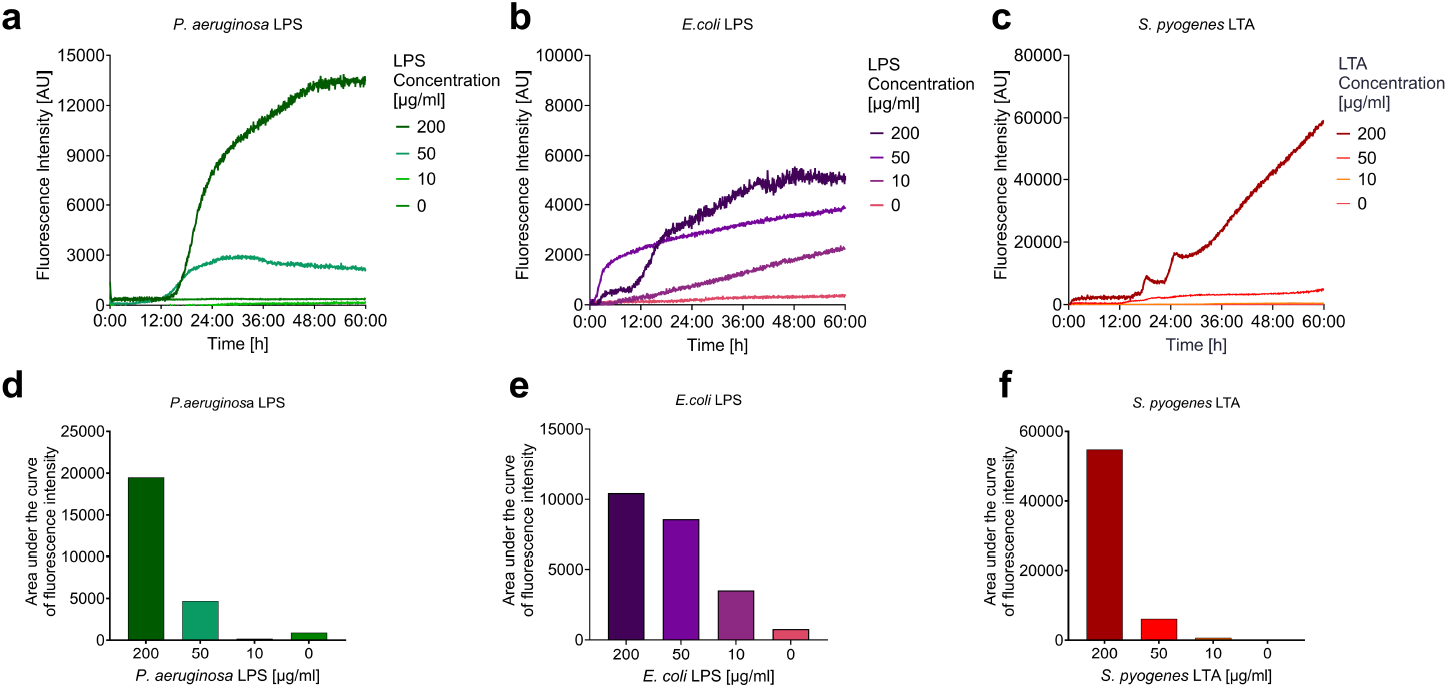
Bacterial surface molecules induce SEVI formation. Freshly dissolved PAP248-286 at a concentration of 400 µg/ml was incubated with indicated concentrations of (a) *P. aeruginosa*-derived LPS, (b) *E. coli*-derived LPS, or (c) *S. pyogenes*-derived LTA at 37 °C in the presence of 25 µM Thioflavin T. Fluorescence intensity was measured every 5 min at an excitation of 450 nm and emission at 490 nm using a Synergy H1 multi-mode reader (BioTek) with the Gen5 software. All values represent fluorescence intensity derived from averaged quadruplicates corrected for the background signal. (d-f) Quantified area under the curve for the data shown in panels a-c. Figure was constructed using GraphPad Prism version 10.

In addition to LPS, we tested the effect of lipoteichoic acid (LTA) from Gram-positive bacteria. Interestingly, a concentration of 200 µg/mL LTA also promoted robust aggregation of PAP248-286 (Fig. 3c). Notably, the extent of fibrillation increased progressively over time, with LTA producing a sustained and significant enhancement in fibril formation comparable to the effects that were observed for LPS. These effects were further quantified through the analysis of the area-under-the-curve (AUC) of the kinetics graphs, which demonstrated that LPS and LTA at high concentrations, and LPS even at low concentrations induce amyloid formation (Fig. 3d-f).

Since contact of PAP248-286 with the bacterial surface molecules LPS and LTA efficiently initiated fibril formation, we sought to determine whether these induced fibrils exhibit antibacterial activity. Using *P. aeruginosa* as a model, we compared the antimicrobial effects of these fibrils to SEVI fibrils generated through mechanical shaking. Notably, both fibril types displayed similar antibacterial activity, supporting the hypothesis that bacterial surface structures like LPS and LTA can induce SEVI fibril formation with properties comparable to those produced by mechanical agitation (Fig. 4).

**Fig. 4.**
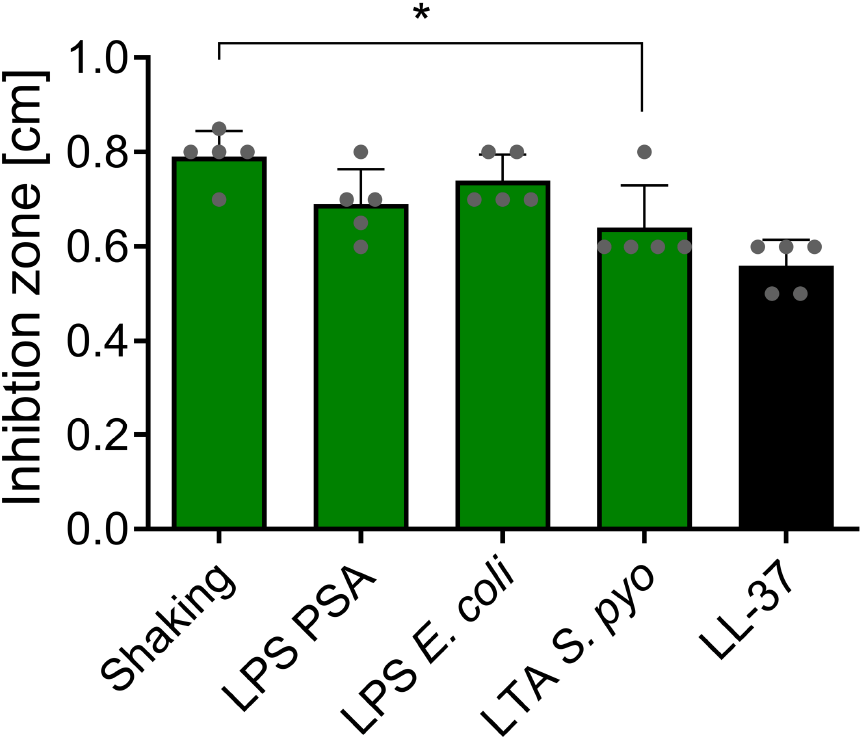
Antimicrobial activity of differently induced SEVI. PAP248-286 was incubated with either lipopolysaccaride (LPS) derived from *Pseudomonas aeruginosa* (LPS PSA), LPS derived from *Escherichia coli* (LPS *E. coli*) or lipoteiconic acid (LTA) derived from *Streptococcus pyogenes* (LTA *S. pyo*). Then antimicrobial activity against *P. aeruginosa* was investigated and compared to SEVI obtained by shaking. Depicted is the mean + standard deviation of five independent experiments. A Mann-Whitney-U test was performed. GraphPad Prism version 10 was used to construct this figure.

### Membrane integrity and intracellular effects of PAP248-286

Many AMPs exert their bactericidal effects by disrupting bacterial membranes. To assess whether non-fibrillar PAP248-286 interferes with bacterial membrane integrity, several assays were conducted. First, the uptake of the fluorescent dye SYTOX, which indicates loss of bacterial membrane integrity, was measured in *L. monocytogenes* and *P. aeruginosa* at pH 7.0 and pH 4.5 (Fig. S3). Measurements for *L. monocytogenes* demonstrated limited membrane damage only at high concentrations of PAP248-286 (Fig. S3a, b). Comparable membrane damage was observed at high PAP248-286 concentrations at pH 7.0 and slightly increased at low pH for *P. aeruginosa* (Fig. S3c, d). To investigate intracellular pH changes, which indicate a potential pore formation, a pHluorin assay was employed using *L. monocytogenes* and *S. agalactiae*, with the AMP Nisin serving as a positive control ^29^. Nisin, PAP248-286 did not induce pore formation in either bacterial species, as indicated by the absence of fluorescence changes associated with pore formation (Fig. S3).

The structural effects of PAP248-286 on bacterial cells were further visualized by electron microscopy. Bacterial cells of *L. monocytogenes* and *P. aeruginosa* were incubated with PAP248-286 at concentrations of 50 µg/mL for one hour. At a concentration of 50 µg/mL morphological changes were observed in *P. aeruginosa* (Fig. 5) and *L. monocytogenes* (Fig. S4), with some bacterial cells showing signs of cellular destruction. Interestingly, intracellular electron-dense clusters were prominent in many *P. aeruginosa* cells treated with 50 µg/mL of PAP248-286 (Fig. 5b). These clusters were further analyzed using energy-dispersive X-ray spectroscopy, which revealed markedly increased phosphate concentrations within these structures, suggesting they are polyphosphate granules (Fig. 5c). These findings suggest a novel intracellular response of *P. aeruginosa* to PAP248-286 that does not rely on membrane disruption but may involve metabolic or stress-related pathways.

**Fig. 5.**
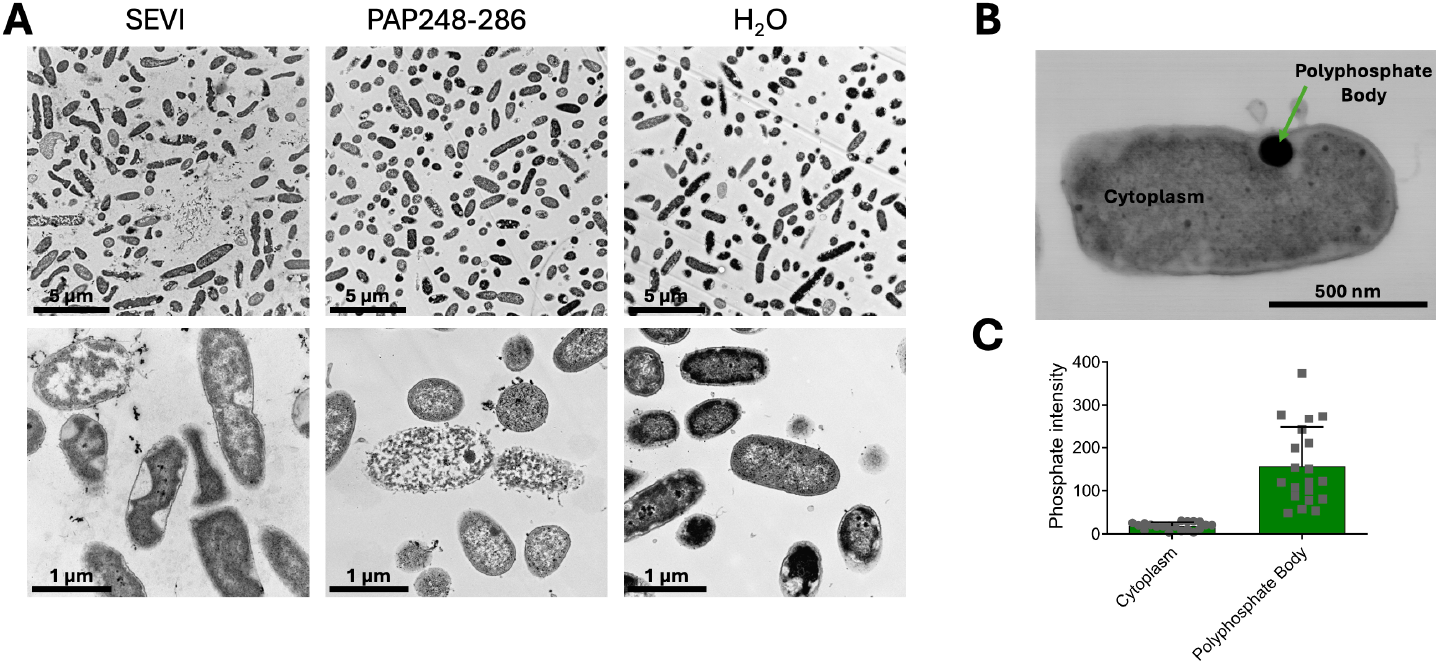
Electron microscopy of *P. aeruginosa* treated with fibrillar and non-fibrillar PAP248-286. Bacterial cells were incubated with 50 µg/ml of fibrillar or non-fibrillar PAP248-286 at pH 4.5. After 60 min of incubation cells were fixed and prepared for electron microscopy. **a)** Images were taken with a transmission electron microscope at different magnifications. **b)** Energy dispersive X-ray spectroscopy of granules and cytoplasm of *P. aeruginosa* treated with non-fibrillar PAP248-286 was performed to measure the elemental composition of granules versus cytoplasm. **c)** 20 cells with polyphosphate bodies were analyzed with EDX and phosphate intensity between the granules and the cytoplasm was compared.

## Discussion

SEVI, an amyloid fibril derived from the PAP248-286 peptide in human seminal plasma, is primarily known for enhancing HIV-1 infection and has also been reported to facilitate bacterial aggregation and phagocytosis ^14^. Our study adds to these findings by providing the first evidence of direct antibacterial activity of non-fibrillar and fibrillar PAP248-286 against clinically relevant bacterial pathogens, including ESKAPE organisms. These findings expand our understanding of the role of SEVI in the mucosal immune defense and highlights its potential as a natural antimicrobial agent.

### Direct antibacterial activity of PAP248-286/SEVI

PAP248-286 in its non-fibrillar form and SEVI fibrils exhibited robust antibacterial activity against a range of clinically relevant pathogens, including ESKAPE organisms and urogenital pathogens such as *S. agalactiae* and *L. monocytogenes*. This activity was dose-dependent and observed under physiologically relevant conditions, including acidic environments resembling those of the vaginal mucosa. Notably, acidic pH enhanced the efficacy of SEVI against certain pathogens, including *P. aeruginosa* and *E. coli*, where bacterial survival was reduced by more than two log levels within two hours at SEVI concentrations comparable to those found in semen. The enhanced activity at acidic pH likely reflects the increased protonation of the amino acid residues of SEVI, such as histidine and aspartic acid, which may optimize electrostatic interactions with negatively charged bacterial membranes ^30 31^. This aligns with findings from other AMPs, whose activity is potentiated in acidic conditions. However, the response to pH was not uniform across bacterial species; for example, activity against *K. quasipneumoniae* was reduced at low pH, highlighting potential species-specific variations in the mechanism of action.

### Role of SEVI fibrils

Our findings demonstrate that the fibrillar structure of SEVI plays an important role in its antibacterial activity. Preincubation with EGCG progressively abolished SEVI’s antimicrobial effects, compatible with the conclusion that fibrils are the primary bioactive form, which is the case for other functions. EGCG is a polyphenol that destroys and remodels SEVI amyloid fibrils in a way that after more than 24h incubation the remaining oligomeric structures are no longer able to seed and induce fibril formation ^32^. However, freshly dissolved PAP248-286, representing the non-fibrillar state of SEVI, also exhibited antibacterial activity. This raises the possibility that the peptide, even in its initial monomeric form, may assemble into antibacterial fibrils over time under the experimental conditions. Notably, factors such as LPS or LTA, which are abundant in the presence of bacteria, can accelerate the fibrillation process. Freshly dissolved PAP248-286 may rapidly transition into its fibrillar state upon interaction with bacterial surface molecules. Such interactions could explain the observed antibacterial effects of non-fibrillar preparations, as they likely contain a mix of monomers, oligomers and fibrils or fibril seeds by the time of testing.

Although SEVI fibrils displayed comparable or superior activity to monomers in radial diffusion assays, their larger size and reduced diffusivity in agar raise questions about how they exert their effects. The fibrillar state may enhance SEVI’s interactions with bacterial surfaces, potentially through multivalent binding or prolonged stability. These properties could compensate for its reduced ability to diffuse within the agar matrix. However, additional studies are needed to fully delineate the relative contributions of monomers, oligomers, and fibrils in SEVI’s antimicrobial activity, since we cannot exclude an alternative mechanism being responsible for the observed antibacterial effects of the non-fibrillar preparations. In mucosal environments, the structural integrity of SEVI fibrils may confer an advantage by resisting proteolytic degradation, thereby prolonging their bioactivity. Fibrils were also found as the bioactive structure promoting enhanced phagocytosis as reported previously^14^, as well as in our experiments. While these findings suggest a multifunctional role for SEVI fibrils in mucosal defense, further investigations are warranted to clarify the dynamics of fibril formation *in vivo* and to determine how environmental factors, such as pH or the presence of bacterial surface molecules, may influence SEVI’s structural state and activity.

### Interactions with bacterial surface molecules

We demonstrated that bacterial surface molecules, such as LPS and LTA, can induce the rapid fibrillation of PAP248-286. These negatively charged molecules not only mimic physiological conditions in the encounter of PAP248-286 with microbial pathogens but also act as catalytic agents for amyloidogenesis. The ability of bacterial components to induce fibril formation highlights a dynamic interplay between SEVI and pathogens, where bacterial surfaces may inadvertently enhance the antimicrobial efficacy of SEVI by accelerating its transition to the fibrillar state. This mechanism adds to the growing body of evidence that amyloid formation can be triggered by diverse environmental factors, including interactions with other amyloidogenic proteins or surface molecules ^33^. The induction of SEVI fibrillation by bacterial LPS and LTA extends this concept and provides a plausible explanation for the possibility of a rapid fibril generation during bacterial infections.

### Mechanism of antibacterial action

Unlike classical AMPs, which often disrupt bacterial membranes, SEVI appears to exert additional antibacterial effects through a distinct mechanism. Membrane integrity assays revealed limited damage, particularly at neutral pH, and electron microscopy did not show any visual indication of massive bacterial cell destructions. Instead, we observed the formation of intracellular electron-dense granules, identified as polyphosphate (polyP) granules via EDX analysis (Fig. 5). PolyP granules are a hallmark of bacterial stress responses, typically associated with starvation or environmental stress, and their formation suggests that PAP248-286 interferes with bacterial metabolism ^34^. PolyP granules represent an energy storage mechanisms, that the cell can use to restore regular metabolism under more favorable conditions ^34^. The generation of polyP granules following treatment may thus signify metabolic stress. Taken together our results are compatible with a limited interference of SEVI with bacterial cell integrity in conjunction with triggering a starvation response, which will ultimately result in bacterial cell death.

### Physiological and immunological implications

The dual functionality of SEVI in bacterial aggregation and direct antibacterial activity underscores its potential role in mucosal immunity. During fertilization, SEVI may contribute to maintaining the sterility of the upper reproductive tract by neutralizing bacterial threats and enhancing their clearance by immune cells. The acidic environment of the vaginal mucosa ^31^ likely synergizes with SEVI’s properties, creating a localized defense mechanism against bacterial colonization. Moreover, the ability of SEVI to target antibiotic-resistant pathogens, including ESKAPE organisms, highlights its potential relevance in combating multidrug-resistant infections. While SEVI’s physiological concentration and activity are optimized for the urogenital tract, its antimicrobial properties may inspire the development of amyloid-based therapeutic strategies for other infections.

## Abbreviations

*A. baumannii*: *Acinetobacter baumannii*
AMP: Antimicrobial peptide
ATCC: American Type Culture Collection
DMSZ: German Collection of Microorganisms and Cell Cultures
*E. coli*: *Escherichia coli*
EDX: Energy-Dispersive X-ray Spectroscopy
*E. faecium*: *Enterococcus faecium*
ESBL: Extended Spectrum Beta-Lactamase
*K. quasipneumoniae*: *Klebsiella quasipneumonie*
*L. monocytogenes*: *Listeria monocytogenes*
LPS: Lipopolysaccharide
LTA: Lipoteichoic Acids
MRSA: Methicillin-resistant *Staphylococcus aureus*
PAP: Prostatic Acid Phosphatase
polyP: Polyphosphate
PSA *P. aeruginosa*: *Pseudomonas aeruginose*
SEVI: Semen Enhancer of Viral Infection
*S. aureus*: *Staphylococcus aureus*
*S. aglactiae*: *Streptococcus agalactiae*
*S. pyogenes, S. pyo*: *Streptococcus pyogenes*
ThT: Thioflavin T
VFS: Vaginal Fluid Simulant
VRE: Vancomycin-reistant enterococci

## Author Contribution

Radial diffusion assays and survival assays with different bacteria were performed and analyzed by SB, EW, NJ, SM and VV. Experiments using EGCG were conducted by FW. Fibril formation was analyzed by LOR. SB, RB and VV did electron microscopy analysis with the support of UR and CR. UR conducted EDX-analysis. Phagocytosis experiments were done and analyzed by US, JZ and PF-P. NJ, SB and FW investigated and evaluated effects on membrane integrity. Experiments were designed by JM and BS. The manuscript was prepared and edited by VV, LOR, JM and BS.

## Acknowledgments

VV, RB, LRO, NJ, UR, CR, JM, and BS thank the German Research Foundation (DFG) for funding within the CRC1279.

## Ethical standards

Experiments were performed in accordance with German laws.

## Competing interests

The authors declare that the research was conducted in the absence of any commercial or financial relationships that could be construed as a potential conflict of interest.

## Data availability

The raw data supporting the conclusions of this article will be made available by the authors, without undue reservation, to any qualified researcher.

## Notes

### Competing Interest Statement

The authors have declared no competing interest.

### Summary of Updates

Figure 1B was updated with new data.

