## Supplemental Figures and Tables for "SEVI Fibrils are Induced by Bacterial Surface Molecules and Exert Antimicrobial Activity Against ESKAPE Pathogens"

Table of Contents

**Figure S1: Influence of fibrillar and non-fibrillar PAP248-286 on phygocytotic uptake by *Staphylococcus aureus*.** **2**

**Figure S2: Activity of non-fibrillar versus fibrillar PAP248-286 against *Pseudomonas aeruginosa*.** **3**

**Figure S3: Bacterial membrane disruption was assessed by monitoring SYTOX and pHluorin fluorescence intensity.** **4**

**Figure S4: Electron microscopy of *Listeria monocytogenes* treated with PAP248-286.** **5**

Table S1: Activity of fibrillar and non-fibrillar PAP248-286 against *Staphylococcus aureus* 6

Table S2: Activity of epigallocatechin gallate (EGCG) against *Pseudomonas aeruginosa* 6

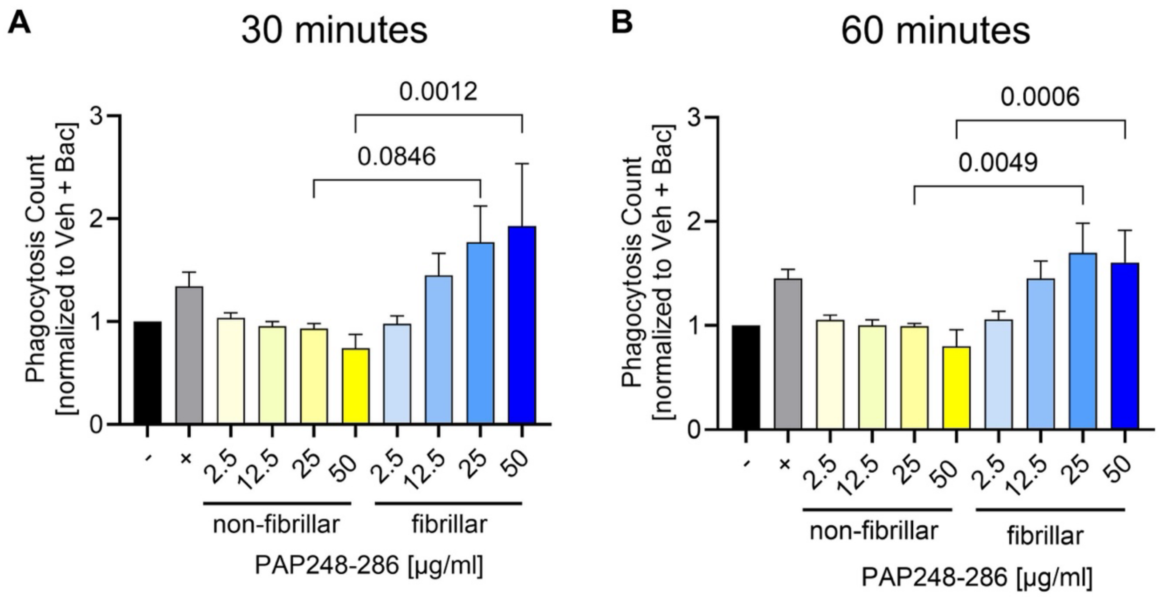


**Fig S1 Influence of fibrillar and non-fibrillar PAP248-286 on phygocytotic uptake by *Staphylococcus aureus***

THP-1 derived macrophages were incubated with pHrodo Red *S. aureus* BioParticles and either fibrillar or non-fibrillar PAP248-286 in concentrations ranging from 2.5-50 µg/ml. An IncuCyte S3 Live Cell Analysis System (Sartorius) took four pictures after (A) 30 min and after (B) 60 min and the number of positive signals was automatically determined using the IncuCyte 2019B Basic Analysis Software module. Significance levels were calculated using a One-way ANOVA.

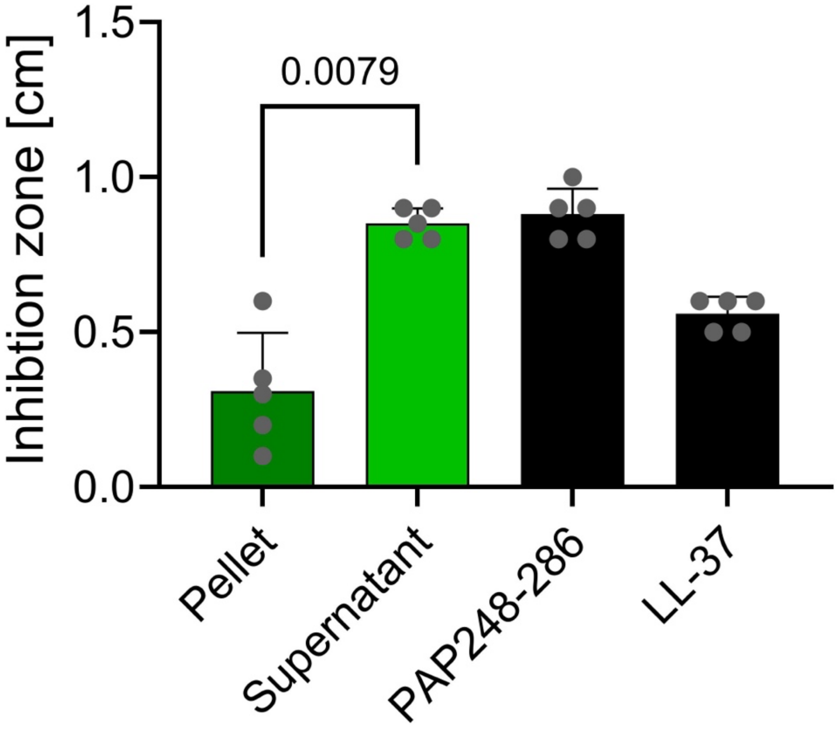


**Fig. S2 Activity of non-fibrillar versus fibrillar PAP248-286 against *Pseudomonas aeruginosa***
Freshly dissolved PAP248-286 was agitated for 17-18 h at 37°C and then centrifuged. In an RDA against *P. aeruginosa,* the inhibitory activity of the pellet, containing high amounts of fibrillar PAP248-286, was compared to the activity of supernatant, which mostly contained the non-fibrillar form. As control feshly dissolved PAP248-286 and LL-37 were additionally tested. Depicted is the mean+ standard deviation of five independent experiments.

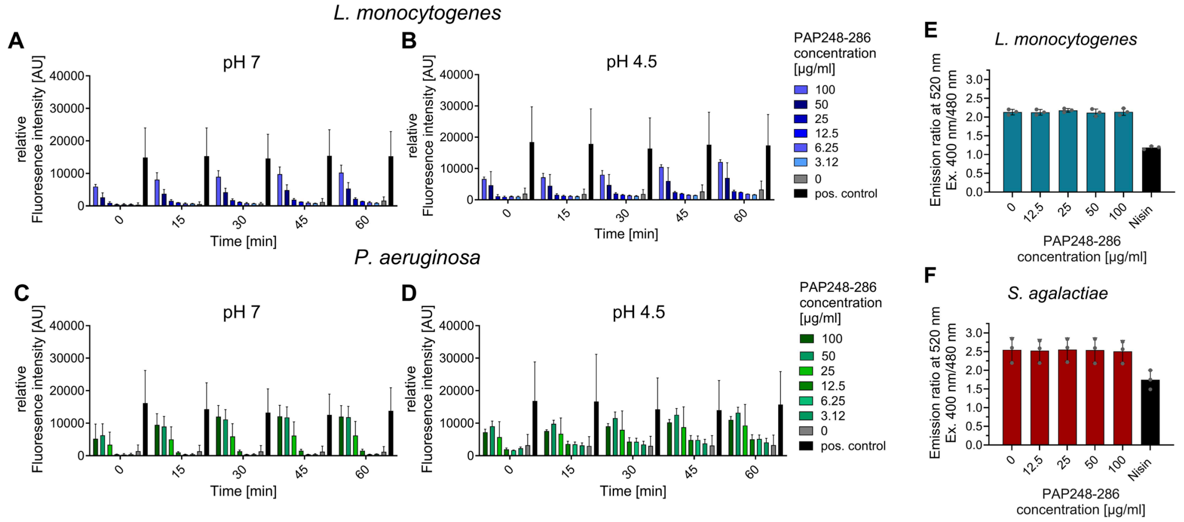


**Fig. S3 Bacterial membrane disruption was assessed by monitoring SYTOX and pHluorin fluorescence intensity**

SYTOX fluorescence intensity (Excitation: 488 nm, Emission: 530 nm) was monitored every 15 min over a time course of 60 min. The bacterial cells were subjected to PAP248-286 concentrations ranging between 100-3.12 µg/ml. Bacteria treated with 70% ethanol served as a positive control in all SYTOX fluorescence experiments. Data represent the mean + standard deviation of three independent experiments. *Listeria monocytogenes* were exposed to PAP248-286 in a neutral pH (A) and in acidic pH (B). In the same line, *Pseudomonas aeruginosa* cells were incubated at neutral (C) and acidic pH (D) with multiple PAP248-286 concentrations. In a phluorin-Assay, (E) *L. monocytogenes* and (F) *Streptococcus agalactiae* cells, carrying pNZ-pHin2$^{LM}$ were treated with fibrillar PAP248-286 at concentrations ranging from 0 to 100 µg/mL in listeria minimal buffer (pH 6.2) for 30 minutes. Fluorescence emission at 520 nm was recorded following excitation at 400 and 480 nm and the respective ratio was calculated. For *L. monocytogenes*, 10 µg/mL nisin was used as a positive control, whereas 100 µg/mL nisin was employed for *S. agalactiae*. Results are shown as the mean ± standard deviation of three independent experiments.

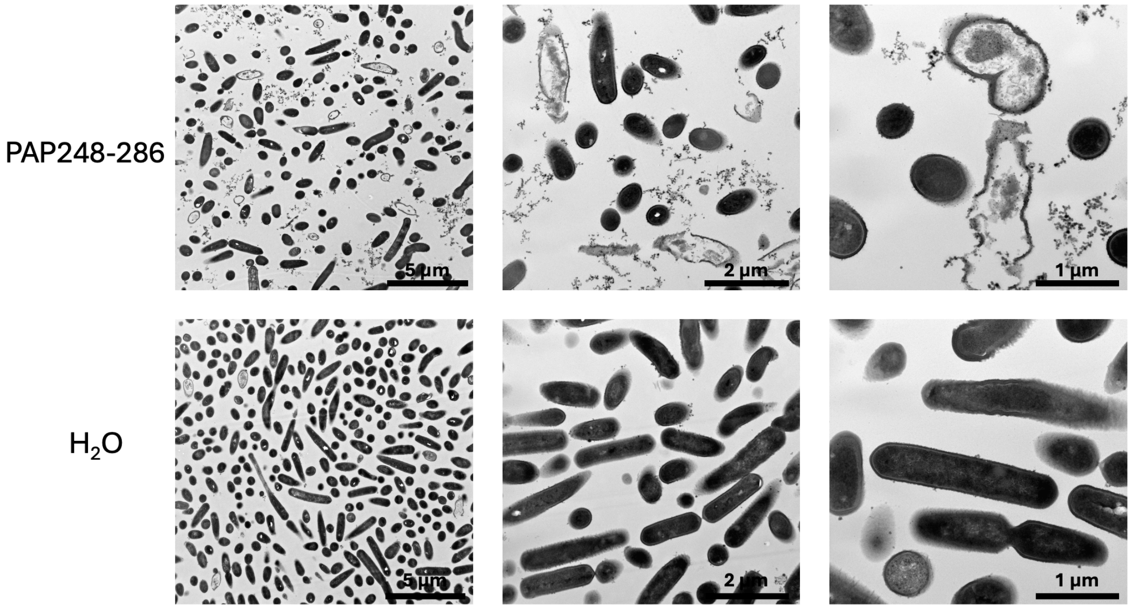


**Fig S4 Electron microscopy of *Listeria monocytogenes* treated with PAP248-286**

*L. monocytogenes* was incubated with non-fibrillar 100 µg/ml PAP248-286 or water (control) for 60 min at a pH of 4.5. Subsequently, cells were fixed and prepared for transmission electron microscopy. Images were taken with a JEOL 1400 Transmission Electron Microscope.

Table S1: Activity of fibrillar and non-fibrillar PAP248-286 against *Staphylococcus aureus* and VRE (n=3)

| **Bacterial species** | **Non-fibrillar PAP248-286 [µg/ml]** | | | | | **Fibrillar PAP248-286 [µg/ml]** | | | | | **Nisin [µg/ml]** |
|---|---|---|---|---|---|---|---|---|---|---|---|
| | **100** | **50** | **25** | **12.5** | **6.25** | **100** | **50** | **25** | **12.5** | **6.25** | **100** |
| ***S. aureus*** | 0 | 0 | 0 | 0 | 0 | 0 | 0 | 0 | 0 | 0 | 1,43 cm |
| **VRE** | 0 | 0 | 0 | 0 | 0 | 0 | 0 | 0 | 0 | 0 | 1,28 |

Table S2: Activity of epigallocatechin gallate (EGCG) against *Pseudomonas aeruginosa* (n=5)

| | **Experiment 1** | **Experiment 2** | **Experiment 3** | **Experiment 4** | **Experiment 5** |
|---|---|---|---|---|---|
| 1mM EGCG | 0 | 0 | 0 | 0 | 0 |
